# scRepertoire 2: Enhanced and Efficient Toolkit for Single-Cell Immune Profiling

**DOI:** 10.1101/2024.12.31.630854

**Authors:** Qile Yang, Ksenia R. Safina, Nicholas Borcherding

**Author notes:** Correspondence: Nicholas Borcherding, 660 S. Euclid Ave, Box 8118, St. Louis, MO 63110.

## Abstract

Single-cell adaptive immune receptor repertoire sequencing (scAIRR-seq) and single-cell RNA sequencing (scRNA-seq) provide a transformative approach to profiling immune responses at unprecedented resolution across diverse pathophysiologic contexts. This work presents scRepertoire 2, a substantial update to our R package for analyzing and visualizing single-cell immune receptor data. This new version introduces an array of features designed to enhance both the depth and breadth of immune receptor analysis, including improved workflows for clonotype tracking, repertoire diversity metrics, and novel visualization modules that facilitate longitudinal and comparative studies. Additionally, scRepertoire 2 offers seamless integration with contemporary single-cell analysis frameworks like Seurat and SingleCellExperiment, allowing users to conduct end-to-end immune profiling with transcriptomic data. Performance optimizations in scRepertoire 2 substantially reduce computational time and memory usage, addressing the demands of the ever-increasing size and scale of single-cell studies. This release marks an advancement in single-cell immunogenomics, equipping researchers with a robust toolset to uncover immune dynamics in health and disease.

## Introduction

High-throughput sequencing technologies are foundational to modern biomedical innovation, driving advancements across various fields. Single-cell technologies stand out for their unparalleled ability to dissect cellular heterogeneity and dynamics across tissues and conditions, making them indispensable in immunology and oncology research. By combining scRNA-seq with scAIRR-seq, derived either through direct sequencing or inference from transcriptomic data, researchers can concurrently analyze gene expression and immune receptor diversity at the single-cell level [1]. The capability to profile both the transcriptional states and clonotype structures of T and B cells through scRNA and scAIRR data provides a robust framework for tracking immune cell activation, clonal expansion, and persistence. These are critical parameters for assessing vaccine efficacy, evaluating immune responses in cancer, responses in cellular therapies, like CAR-T, and elucidating mechanisms underlying autoimmune diseases [2].

Despite the rapid growth of scAIRR analysis, limitations remain with existing tools, primarily available in Python and R [3]. Many tools lack robust integration for immune receptor profiling with transcriptomic data or flexibility in data export formats, which hinders reproducibility and cross-platform compatibility in expanding datasets. The increasing demand for standardized, flexible, and scalable analysis frameworks underscores the need for improved software solutions in this field. The growing size and scope of single-cell data creation also underscores this need for improved standardization and efficiency.

The initial release of scRepertoire addressed some of these needs, establishing itself as an integral tool within single-cell workflows due to its compatibility with widely used R-based single-cell analysis platforms like Seurat and SingleCellExperiment [4]. scRepertoire enabled researchers to merge, filter, and visualize clonotype data, offering essential resources for the indepth characterization of immune repertoires at the single-cell level. As one of the first open-source packages for scAIRR analysis, scRepertoire has over 27,000 downloads from Bioconductor and is a broad utility for the field. Nevertheless, much like the fast-paced changes in scRNA analysis, scRepertoire required a significant rebuild to support new and emerging analyses.

Here, we present the latest release of scRepertoire, which introduces a suite of powerful new features and enhancements aimed at refining immune receptor analysis and visualization. This update includes advanced tools for summarizing amino acid sequences, such as positional entropy and residue-specific composition profiles, alongside visualizations of k-mer distributions and V-J pairings. User utilities have been optimized for speed, with faster generation of clonal pairs, improved clustering, and expanded support for importing and exporting clone data across data types. Additional features include identifying T and B cell receptor (TCR/BCR) doublets and improved repertoire comparisons with customizable normalization. scRepertoire seamlessly interacts with deep learning modules such as Trex, Ibex, and ImmApex and other packages, such as APackOfTheClones, scPlotter, and DandelionR, facilitating a more comprehensive and scalable approach to single-cell immune profiling [5–9].

## Design and Implementation

### Workflow

The scRepertoire package provides a comprehensive, R-based framework for immune repertoire analysis, seamlessly integrating clonotype data with transcriptomic profiles to enable sophisticated insights into immune cell populations (Figure 1). The recent redesign of scRepertoire emphasizes accessibility and usability, with universalized function names and language to make the package approachable for novice and experienced users. Documentation has been significantly expanded, and a dedicated pkgdown website now hosts extensive examples and tutorials, creating a readily accessible knowledge base for users to maximize the package’s capabilities.

**Figure 1:**
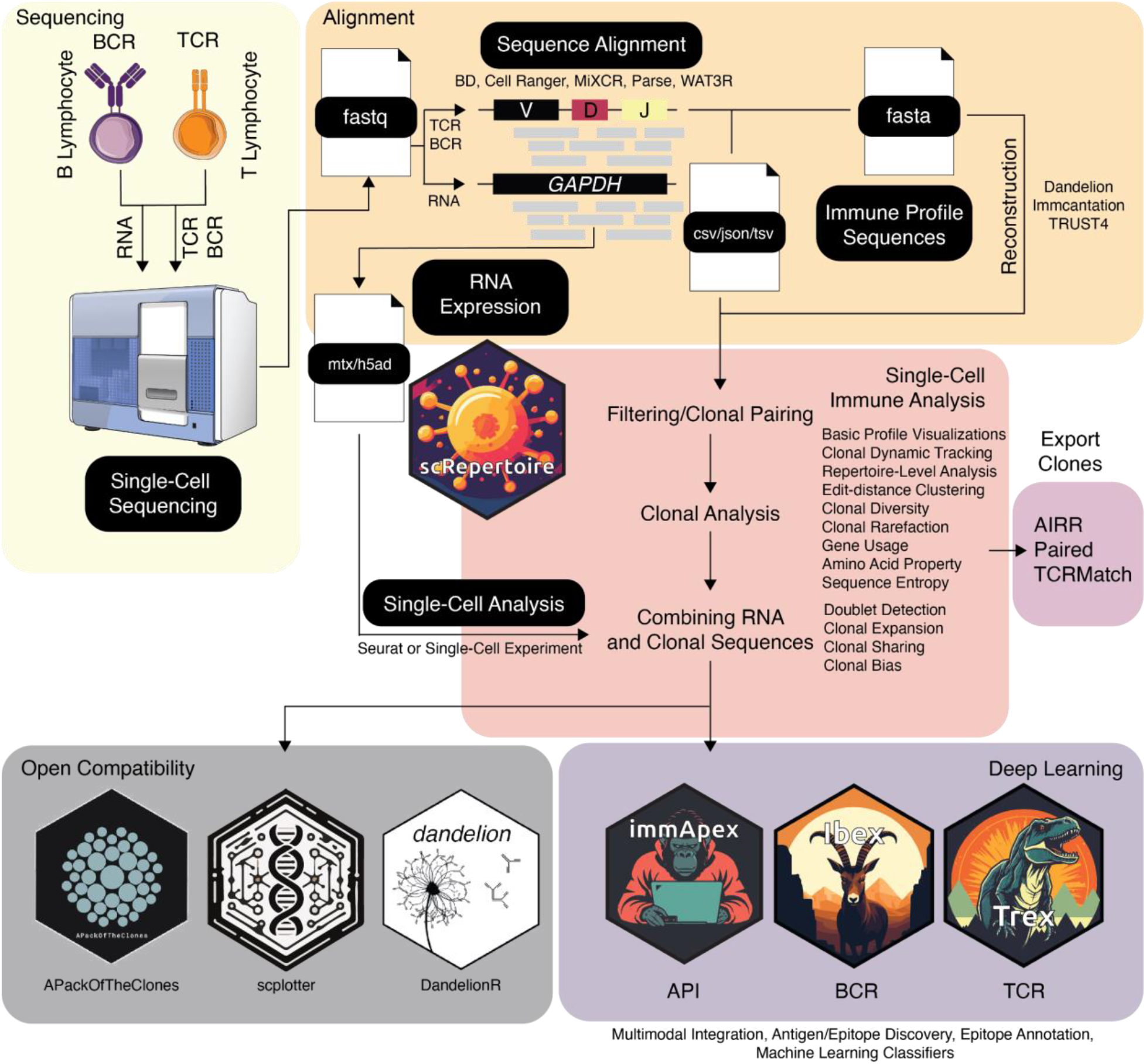
Schematic representation of the scRepertoire workflow and analysis. scRepertoire workflow overview with immune profiling sequence alignments utilized directly from common pipelines or reconstruction and loaded into the scRepertoire environment. scRepertoire will perform filtering and pairing of the respective adaptive immune receptor and offer a large array of analyses and visualizations. The resulting clonal pairs can be attached to single-cell RNA/protein/chromatin data processed via Seurat or SingleCell Experiment pipelines. scRepertoire has native compatibility with immApex, Trex, and Ibex, allowing for the deep-learning-based embedding of the scAIRR sequences for multimodal integration or classifier building.

This workflow processes data from multiple sequencing and alignment formats (e.g., TSV, JSON, CSV) and is compatible with outputs from widely used alignment pipelines, supporting flexibility in data input. Following data import, scRepertoire performs stringent clonal pairing and quality control to assign clonotypes at the single-cell level. The assignment of paired clones allows for robust clonal analysis that quantifies and visualizes clonal expansion and lineage relationships within B-cell and T-cell populations. By linking RNA expression data with clonal sequences, scRepertoire provides an integrative view of immune clonotypes within their transcriptional context, which is essential for uncovering immune cells’ functional states and diversity. In addition to core analysis capabilities, scRepertoire supports downstream applications through compatible tools: Trex, Ibex, and immApex, which aid in the development of models and classifiers for the TCR, BCR, or adaptive immune receptors, respectively. These extensions allow users to tailor their analyses toward specific receptor types and leverage advanced computational techniques, including machine learning, for predictive modeling.

### Expanded Data Compatibility

scRepertoire 2 supports a range of scAIRR-seq formats, including 10x Genomics, AIRR, BD Rhapsody, MiXCR [10], Omniscope, Parse Bio Evercode, TRUST4 [11], and WAT3R [12]. With the additional format support, a new data importer `loadContigs()`has been introduced and automatically detects input format. For exporting data, we added `exportClone()`, a flexible solution for exporting scRepertoire clonal data, making it compatible with multiple formats for easy integration into downstream analysis tools. This function allows users to save tabular clone information organized by barcodes and sequence details such as those compatible with AIRR, supporting seamless data sharing and interoperability across pipelines. In addition, scRepertoire now allows custom clonal definitions in a cell-wise metadata column so that clones identified with alternative pipelines can be visualized/analyzed within scRepertoire.

### Performance Optimizations

With the integration of C++ source code via Rcpp, essential methods have been re-engineered internally to significantly reduce both practical runtime overhead and theoretical time complexity. The most important enhanced methods include those for generating clonal pairs for both with `combineTCR()` and `combineBCR()`, where the runtime of the receptor sequence - cell pairing step now scales linearly instead of quadratically with respect to repertoire size. For BCR clonotype inference and general clonal clustering based on Levenshtein distance, the runtime has also been reduced to scale linearly. In addition, threshold length filtering has been implemented when clustering clones by edit distance. A similar optimization was performed to calculate amino acid and nucleotide-based Kmer counts. For all visualization functions, we implemented a conditional evaluation to prioritize the order of operations based on the user’s specified output. Specifically, if the user selects data export and plotting functionalities, the package now evaluates and executes data export tasks before initiating plotting.

### Enhanced Repertoire Summarization

The latest release of *scRepertoire* introduces advanced features for comprehensive immune repertoire summarization, focusing on amino acid composition and V(D)J gene usage. The `positionalProperty()` function facilitates the examination of physical properties along the CDR3 sequence, enabling detailed analysis of specific sequence regions potentially involved in antigen specificity or structural stability. Additionally, the `positionalEntropy()` function quantifies variability along CDR3 sequences by measuring entropy at each residue, allowing for identifying conserved or highly variable motifs that may have implications for epitope recognition. Further augmenting these capabilities, the suite of amino acid compositional tools enhances the analysis of amino acid and sequence motifs. The `percentAA()` function calculates amino acid composition frequencies across receptor sequences, supporting the detection of conserved amino acid signatures across clonotypes. Similarly, `percentKmer()` evaluates k-mer distribution, providing a granular assessment of recurring sequence motifs across diverse clonotypes or experimental conditions. Finally, `percentVJ()` visualizes the frequency and distribution of V and J gene pairings in an intuitive heatmap format, facilitating the identification of overrepresented gene combinations within immune repertoires.

### Clonal Diversity Analysis

The `clonalRarefaction()` function in scRepertoire offers a versatile framework for rarefaction analysis, allowing users to estimate clonal richness while accounting for potential sampling biases. This capability is especially valuable in comparative studies of immune responses across diverse experimental conditions, where controlled adjustments for sampling depth are essential to ensure accurate cross-sample comparisons. The function’s grouping feature further enhances its utility by enabling structured comparisons across experimental or biological conditions, thereby supporting a robust and nuanced assessment of clonal diversity under varied settings. Additionally, scRepertoire has streamlined the `StartracDiversity()` function, removing custom data dependencies to enhance computational efficiency and accessibility. This re-implementation facilitates estimating differential clonal expansion, cross-tissue migration, and state transitions, offering critical insights into clonal dynamics within and between tissue environments.

### Empowering ML Applications

Building on the established Seurat and Bioconductor SingleCellExperiment infrastructure, our VDJ integration builds a direct bridge to structured data for machine/deep learning. An application example built upon scRepertoire 2 is the amino acid autoencoders Trex [5] and Ibex, which can leverage scAIRR sequences to create meaningful latent dimensions. These latent dimensions can be used independently or incorporated into multimodal single-cell analysis workflows. Both models use ImmApex [9], a scRepertoire-compatible companion toolkit that facilitates deep and machine learning for immune receptor sequences for classification or predictive modeling.

## Results

An example data set for exploring scRepertoire capabilities is provided by Jiang et al. [13]. This data set includes single-cell RNA and immune profiling (both TCR and BCR) of erythema migrans lesions (SKL) and adjacent normal skin (SKN), offering a unique view into immune activity in distinct tissue microenvironments. The cohort consists of six distinct patients. However, two samples were removed for the following analyses based on the lack of scAIRR sequences or lack of sequencing of a secondary tissue. After quality control filtering, individual lymphocytes with productive scAIRR sequences were retained, identifying 13 distinct cell types (Figure 2A). These cell types demonstrated varied clonal expansion profiles, providing insights into immune cell dynamics within the tissue (Figure 2B).

**Figure 2:**
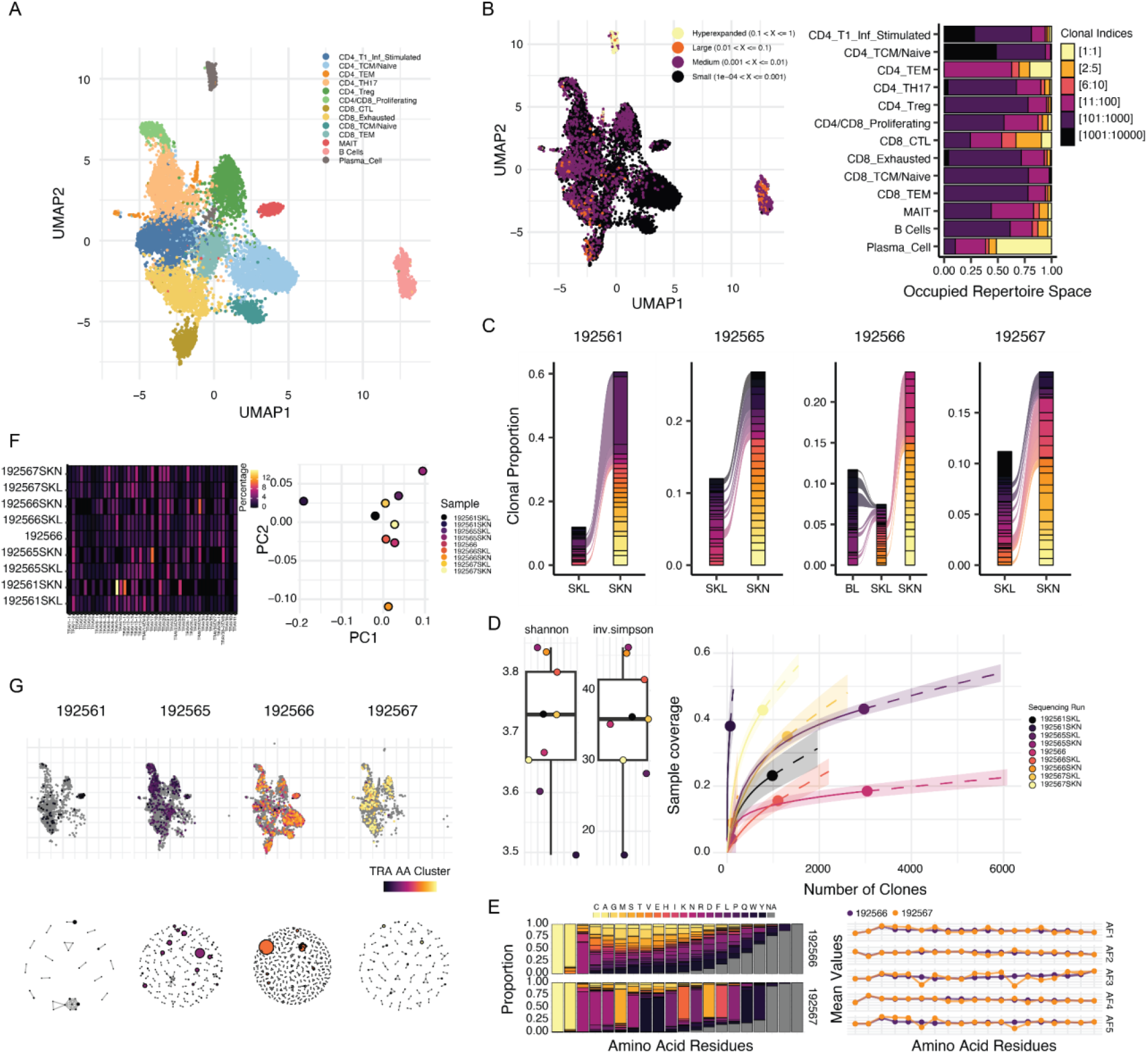
Single-cell analysis of lymphocytes in erythema migrans lesion and adjacent normal skin. **A**. UMAP projection showing lymphocyte subpopulations identified from single-cell RNA and TCR/BCR sequencing data. **B**. UMAP overlay highlights clonal expansion by relative proportion, demonstrating the distribution of expanded clones and bar plots representing occupied clonal space, ranked by clone size and stratified by lymphocyte cell type. **C**. Alluvial plots depicting TCR clones sharing between erythema migrans lesions (SKL), adjacent normal skin (SKN), and blood (BL), highlighting the top 20 clones for each patient by their repertoire proportion. **D**. Bootstrapped Shannon and inverse Simpson diversity values are plotted for each sequencing run to assess repertoire richness with clonal rarefaction and extrapolation analysis estimating sample completeness through Shannon diversity indices. **E**. Percentage of amino acid usage and mean Atchley factor values along the CDR3 amino acid sequences for the heavy chain for 192566 and 192567 samples.. **F**. Heatmap of TRAV gene usage with a principal component analysis across sequencing runs. **G**. TRA chain clustering based on amino acid sequences using a normalized Levenshtein edit distance threshold of 0.85 overlaid onto a UMAP and using a Fruchterman-Reingold layout in igraph. The individual dot size denotes the number of sequences or clonal expansion.

Plasma cells exhibited the most prominent clonal expansion among the identified cell types. In the 192567SKL sample, a single clone accounted for over 50% of the cluster, highlighting its dominant presence (Figure 2B). Other cell types with significant expansion included CD4+ tissue effector memory (CD4_TEM) cells, cytotoxic CD8+ T (CD8_CTL) cells, and mucosal-associated invariant T (MAIT) cells. These findings underscore the diversity of clonal activity across immune cell populations.

To further explore immune dynamics, the top 20 TCR clones were examined for clonal sharing across tissues within the same patient (Figure 2C). Notably, overlap was observed between SKN and SKL samples, suggesting potential roles for tissue-resident T cells in both environments. In the 192566 sample, which included sequencing from peripheral blood (BL), TCR clones shared between BL and SKL samples were mutually exclusive of SKN samples. This finding suggests specific trafficking or retention mechanisms. Interestingly, no BCR clonal sharing was observed across tissues, potentially reflecting different modes of immune compartmentalization or low overall sampling (data not shown).

To quantify repertoire diversity, scRepertoire applied Shannon diversity and inverse Simpson diversity estimates (Figure 2D). By bootstrapping across the smallest repertoire in the comparison group and averaging results from 100 runs, scRepertoire provided robust diversity estimates. Additionally, diversity metrics enabled interpolation and extrapolation using the iNEXT package [14], facilitating rarefaction and sample completeness assessments. These analyses revealed differences in diversity that may correspond to distinct immune pressures or clonal selection processes.

Repertoire summarization within scRepertoire provided additional insights into amino acid use and sequence properties. For the two samples with BCR sequences within multiple tissues, CDR3 sequence analysis revealed a predominant usage of serine (S) at positions 6 and 11 in sample 192567 (Figure 2E). Analysis of amino acid properties, such as Atchley factors [15], indicated positional decreases in molecular size (AF3) and electrostatic charge (AF5) associated with these serine-enriched positions, potentially influencing antigen-binding characteristics.

Additional summarization approaches included calculating the percentage of amino acid k-mers, VJ gene pairing, and single gene usage within scAIRR chains (Figure 2F). These metrics provided a comprehensive overview of repertoire composition. Dimensional reduction techniques, such as principal component analysis, were applied to exported data (Figure 2F).Clustering of clones by nucleotide or amino acid sequence, based on normalized Levenshtein distance (set to 0.85), can enable the identification of potential antigen-driven selection pressures (Figure 2G). Clustering could be performed across all sequences or grouped by categorical variables, such as patient samples. Moreover, scRepertoire supports exporting clustering networks as igraph objects [16], facilitating integration with network-science-based analytical tools.

These analyses illustrate the breadth of scRepertoire’s capabilities for immune repertoire analysis. By integrating sequence-level insights with functional and diversity metrics, scRepertoire provides a robust framework for studying immune responses in complex tissue environments.

## Availability and future directions

scRepertoire is built within the R framework and is freely available under the MIT license. It is hosted on Bioconductor (https://bioconductor.org/packages/scRepertoire). Comprehensive documentation, tutorials, and source code are provided to support users at https://github.com/ncborcherding/scRepertoire. The code and detailed methods for reproducing the erythema migrans data set analysis are available at https://github.com/ncborcherding/scRepertoire.v2_manuscript.

Future development plans for scRepertoire focus on enhancing its interoperability with other immune receptor analysis tools. Improved compatibility will encourage custom extensions and collaborative tool development, fostering the growth of related packages such as APackOfTheClones and DandelionR. These integrations aim to streamline workflows for users analyzing complex immune repertoires.

Another key development area is incorporating advanced statistical methods and machine learning applications within scRepertoire and its compatible extensions. These enhancements will allow the uncovering of deeper insights into immune repertoire dynamics, predict immune responses, and identify biomarkers for clinical and research purposes. In the long term, scRepertoire’s framework should be extended to accommodate multi-omics integration. For example, combining immune repertoire data with spatial transcriptomics or proteomics could provide a more comprehensive picture of immune responses.

## Acknowledgments

From the time of original publishing to scRepertoire 2, the community that uses scRepertoire has contributed consistently to improving the package. We would like to specifically acknowledge the users that directly contributed code to the package: Massimo Andreatta, Nicholas Bormann, Scott Nicholas Furlan, Gloria Kraus, I-Hsuan Lin, Jacqueline HY Siu, Kelvin Tuong, and Panwen Wang. Also, we thank GitHub users Simon-Leonard and Liripo. In addition, we would like to thank Jonathan Noonan, Kyle Romine, and Kelvin Tuong for their constructive suggestions for scRepertoire improvements.

